# O-antigen promotes age-dependent pathogen competition with the gut microbiota and complement evasion during natural systemic dissemination

**DOI:** 10.64898/2026.11.13.766611

**Authors:** Aidan J. Schmidt, Erick D. Nilsen, Marienela Y. Heredia, Victoria G. Zalutskiy, Andres Mejia, Gustavo Caballero-Flores

**Affiliations:** Department of Medical Microbiology & Immunology, University of Wisconsin-Madison, Madison, Wisconsin, 53706, USA; Comparative Pathology Laboratory, University of Wisconsin-Madison, Madison, Wisconsin 53706, USA

## Abstract

Bacterial pathogens employ multiple virulence factors to outcompete the gut microbiota and immune system during host colonization. Among these, O-antigen (O-ag) is a common cell-surface virulence factor in gram-negative bacteria extensively studied *in vitro* and in models of systemic infection. Yet, its role during bacterial enteric infection is less understood. Particularly, O-ag contributions to pathogen competition with the gut microbiota and natural establishment of diarrhea-associated bacteremia remain unclear. Here, we investigate the role of O-ag in pathogenesis of *Citrobacter rodentium*, a murine enteric pathogen widely used as a surrogate model of the human-restricted pathogen Enteropathogenic *Escherichia coli* (EPEC). Using conventional mice of different ages and gnotobiotic mice harboring age-distinct murine microbiotas infected with WT or O-ag-deficient *C. rodentium*, we demonstrate that O-ag is required for pathogen competition with the adult (but not juvenile) microbiota. This age- and microbiota-dependent restriction of the O-ag mutant *in vivo* is, at least in part, metabolite-dependent because cell-free cecal extracts from adults, but not juveniles, inhibit mutant growth, whereas neither extract affects WT growth. Despite effective gut colonization, the O-ag mutant does not elicit significant intestinal inflammation, systemic dissemination, or lethality in immunocompetent juvenile mice. In complement-deficient juveniles, however, the mutant exhibits systemic dissemination comparable to the WT, resulting in substantial mortality. Accordingly, the O-ag mutant displays increased complement deposition in the systemic environment than the WT. Our results support a model in which O-ag is dispensable for colonization of the infant gut but essential for resistance to complement during subsequent establishment of diarrhea-associated bacteremia.

**IMPORTANCE:** The infant and adult gut represent strikingly distinct environments due to differences in microbiota composition, immune development, and epithelial physiology. Yet, whether bacterial pathogens employ conserved or distinct virulence strategies to colonize the infant versus adult gut is unclear. Similarly, the virulence determinants that enable bacterial pathogens, typically considered intestinal residents, to disseminate beyond the gut in susceptible hosts remain poorly understood. This is particularly relevant to EPEC, which can cause severe diarrhea and disseminate systemically in susceptible children but is rarely associated with disease in healthy adults. Here, we show that O-ag is required for colonization of the adult (but not the young) gut and for the establishment of diarrhea-associated bacteremia. Mechanistically, O-ag promotes resistance to microbiota-derived inhibitory compounds during gut colonization and to complement-mediated killing during natural systemic dissemination. Identifying host context-dependent virulence factors may inform tailored therapies for infectious diarrhea and associated complications in vulnerable populations.

## BACKGROUND

Children under age five are at higher risk of developing life-threatening enteric bacterial infections than adults (1). This is partially due to their developing gut microbiota and immune system compared to adults, resulting in impaired colonization resistance to invading pathogens (2,3). This age-dependent phenotype is recapitulated in the *Citrobacter rodentium*-mouse model of infectious diarrhea (ID), in which juvenile mice exhibit enhanced pathogen colonization, disease severity, and lethality compared to adult animals (4,5). *C. rodentium* is a surrogate model for the closely-related, human-restricted enteropathogenic *Escherichia coli* (EPEC) that can cause life-threatening ID in children, but rarely causes disease in healthy adults (6-8). However, because most *C. rodentium* studies have been conducted in adult mice, the mechanisms by which this pathogen, and consequently EPEC, colonizes the infant gut remain elusive.

O-antigen (O-ag) constitutes the highly variable, outermost layer of bacterial lipopolysaccharide (LPS) in the membrane of Gram-negative bacteria (9). LPS and O-ag have been extensively studied *in vitro* and in models of systemic infection, showing they enable pathogen evasion of innate immune factors, including complement (10-12). However, the contributions of O-ag during intestinal bacterial pathogenesis, including its role in pathogen competition with the infant and adult gut microbiota, bacterial translocation across the gut, and natural establishment of ID-associated bacteremia, remain poorly understood.

Leveraging *C. rodentium* infections in juvenile and adult mice, we reveal a previously unrecognized host age-dependency of O-ag during enteric colonization and competition with the gut microbiota. We show that O-ag-deficient *C. rodentium* can successfully colonize the gut of immunocompetent juvenile mice, but can only transition from enteric to systemic infection in complement-deficient juveniles, demonstrating O-ag is dispensable for gut colonization but essential for pathogen systemic dissemination in the presence of an intact immune system. Thus, we demonstrate host-context dependent effects of O-ag during establishment of ID and ID-associated bacteremia in early life. These studies may constitute a foundation for future development of age-tailored preventive therapeutics against ID and its complications.

## RESULTS

### *C.rodentium* O-ag is required for pathogen competition with the adult, but not juvenile, gut microbiota

We previously reported transposon sequencing screens in *C. rodentium* during enteric infection in Specific-pathogen-free (SPF) and Germ-free (GF) adult mice (13), which suggested LPS/O-ag is required for competition with the adult gut microbiota. To validate these observations, we generated a *C. rodentium* Δ*rfaL::Chl* (Δ*rfaL*) mutant lacking RfaL, an O-ag ligase also known as WaaL that binds fully-formed O-ag to LPS (Fig.S1A) (14). As expected, the Δ*rfaL* mutant retained intact Lipid A and core oligosaccharide but lacked O-ag attached to LPS (Fig. S1B). The mutant did not show major growth defects and exhibited robust susceptibility to complement-enriched serum *in vitro* (Fig. S1C,D). Chromosomal transcomplementation of *rfaL* in Δ*rfaL* restored O-ag structure and complement resistance (Fig. S1B-D). These results confirmed Δ*rfaL* is an O-ag-deficient strain.

To assess the role of RfaL and O-ag on *C. rodentium* enteric pathogenesis, we orally gavaged a 1:1 mixture of WT and Δ*rfaL* into adult SPF mice. During co-infection, the WT strain displayed typical colonization kinetics in the adult mouse gut, while Δ*rfaL* was unable to colonize (Fig. 1A). To assess whether Δ*rfaL* colonization defect was microbiota-dependent, we performed WT/Δ*rfaL* co-infections in adult SPF mice with antibiotic-depleted gut microbiota, and found Δ*rfaL* maintained colonization over time, although at lower levels than the WT (Fig. 1B). These results suggest that the O-ag mutant cannot compete effectively with the adult gut microbiota but can successfully colonize mice with impaired microbiota function. Thus, we tested the ability of Δ*rfaL* to colonize juvenile (18-day-old) mice, which exhibit impaired microbiota-mediated colonization resistance and lethal *C. rodentium* enteric infection (4,5). WT/Δ*rfaL* co-infected juveniles resembled microbiome-depleted adult mice, with the mutant colonizing at stable but lower levels compared to the WT (Fig. 1C). We next evaluated the ability of the O-ag mutant to colonize juvenile and adult mice during single infection. Again, Δ*rfaL* failed to colonize the adult gut (Fig. 1D), a defect that was rescued with transcomplementation of *rfaL* (Fig. S1E). In contrast, juveniles were fully permissive to Δ*rfaL* single infection, with colonization kinetics comparable to WT in adult mice (Fig. 1D vs A).

**Figure 1.**
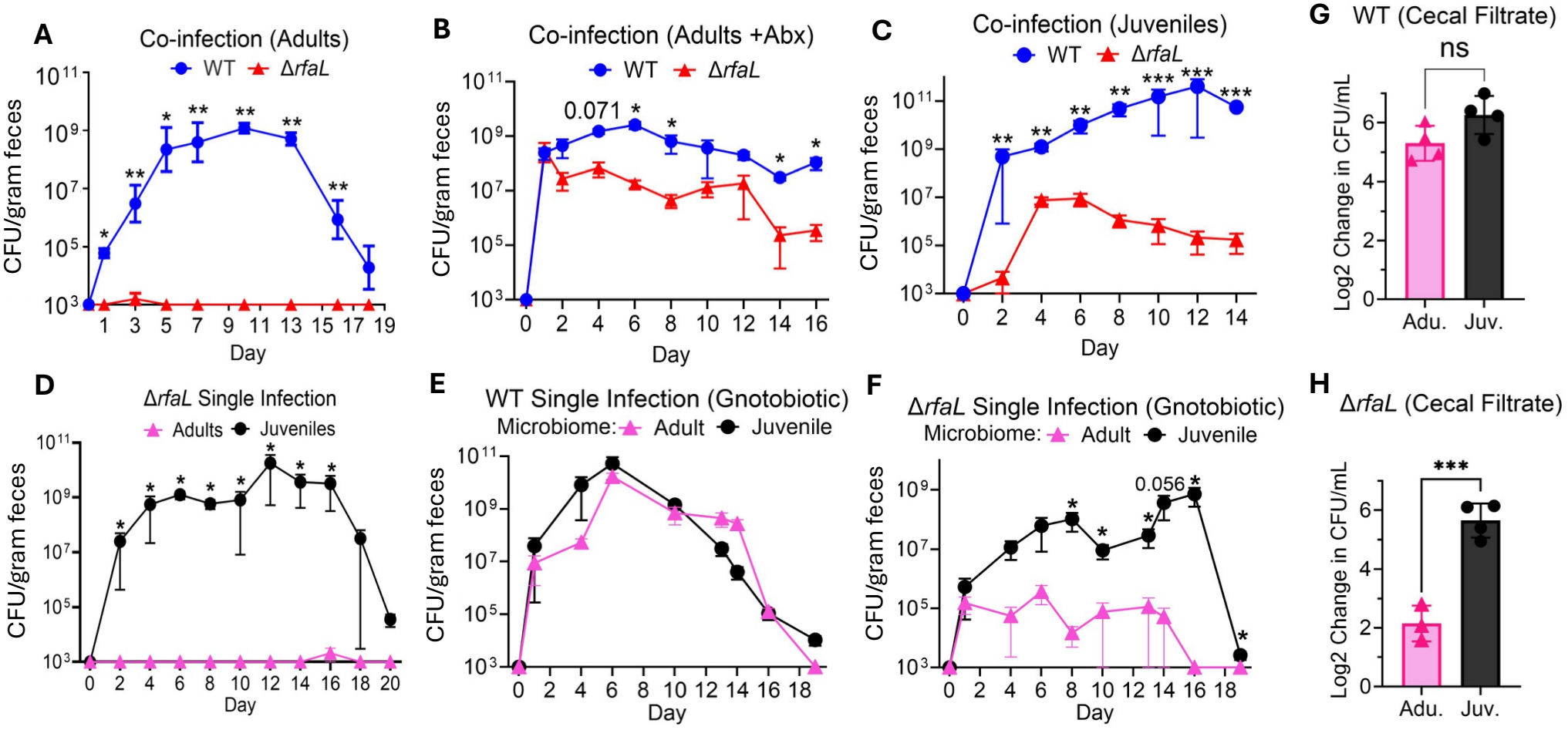
O-ag is required for *C. rodentium* colonization in the presence of an adult, but not juvenile, mouse gut microbiota. **A –** *C. rodentium* wild-type (WT) and Δ*rfaL* co-infection (1:1 ratio) in adult SPF mice. Colony-forming-units (CFU) in feces over the course of infection are shown. *C. rodentium* WT displayed typical, self-limiting colonization kinetics. Δ*rfaL* was unable to colonize the adult mouse gut. **B –** WT and Δ*rfaL* co-infection in adult SPF mice treated with an antibiotic cocktail (vancomycin, neomycin, metronidazole) during infection. CFU in feces are shown. *C. rodentium* WT maintained high colonization levels throughout the experiment. Δ*rfaL* showed a reduced but stable colonization. **C –** WT and Δ*rfaL* co-infection in juvenile SPF mice. CFU in feces during infection are shown. *C. rodentium* WT showed quick, high, and sustained colonization. Δ*rfaL* displayed delayed and low but stable colonization. Juveniles succumbed to infection after day 14. **D –** Δ*rfaL* single infection in adult and juvenile SPF mice. CFU in feces during infection are shown. Δ*rfaL* did not colonize the adult mouse gut during single infection. Δ*rfaL* displayed a self-limiting colonization kinetics in juveniles, typical of the WT in adults (A). **E –** WT single infection in adult gnotobiotic mice harboring adult or juvenile mouse gut microbiotas. No significant differences in colonization were observed. **F –** Δ*rfaL* single infection in adult gnotobiotic mice harboring adult or juvenile mouse gut microbiotas. Δ*rfaL* displayed increased colonization with the juvenile gut microbiota than with the adult type. **G, H –** WT (G) and Δ*rfaL* (H) were cultured anaerobically in cell-free, cecal extract filtrates from adult or juvenile SPF mice. Bars indicate mean Log2 change in CFU after 24 hours compared to time 0. Data are shown as mean ± SEM (A-H). Statistical analyses: Mann-Whitney U test (A-H). **\*** p≤0.05 **\*\***p≤0.01 **\*\*\***p≤0.001. †, Lethality.

To confirm the contribution of the gut microbiota to the age-dependent colonization of the O-ag mutant, we performed cecal microbiota transfers (CMTs) from adult or juvenile SPF donors into adult GF mice, followed by infection with Δ*rfaL* or WT. The microbiota of recipients after CMTs resembled that of donors (Fig. S2). The WT strain displayed no significant differences in colonization of gnotobiotic mice harboring adult or juvenile microbiota (Fig. 1E). In contrast, Δ*rfaL* efficiently colonized mice harboring juvenile microbiota, whereas its levels remained low and near the limit of detection in mice harboring adult microbiota (Fig. 1F). To determine whether microbiota-derived metabolites contribute to the age-dependent inhibition of Δ*rfaL*, we measured *C. rodentium* growth in cell-free cecal extracts from adult and juvenile naive mice. While the WT showed comparable growth in adult and juvenile cecal extracts, Δ*rfaL* displayed impaired growth in the adult, but not juvenile, extracts (Fig. 1G,H). These results indicate that microbiota-derived metabolites contribute to the age-dependent inhibition of the O-ag mutant.

### O-ag is essential for natural establishment of ID-associated bacteremia and complement resistance in juveniles

O-ag is well-known to aid bacterial evasion of complement-mediated killing *in vitro* and in animal models where pathogens are systemically administered (10,12,15,16). Additionally, classic and recent studies have shown that complement-deficient (C3^-/-^) juvenile mice are highly susceptible to enteric infection with *C. rodentium* WT, showing increased pathogen gut colonization and lethality compared with C3^+/+^ counterparts (17-19). To assess whether complement-deficient juveniles are also more susceptible to enteric infection with O-ag-deficient *C. rodentium*, we infected C57BL/6J WT (B6) and C3^-/-^ mice with Δ*rfaL*. There were no significant differences in mutant colonization between mouse genotypes (Fig. 2A), however, substantial mortality in C3^-/-^ compared to B6 juveniles was observed (Fig. 2B). Notably, the lethality caused by the O-ag mutant in C3^-/-^ mice was not accompanied by marked intestinal inflammation (fecal Lipocalin-2) (Fig. 2C) or histological pathology (Fig.2D,S3). Rather, these parameters remained lower in both C3^-/-^ and B6 juveniles infected with Δ*rfaL* than in mice infected with the WT strain. These results suggest that lethality caused by Δ*rfaL* in C3^-/-^ juveniles is not due to enhanced enteric disease severity, but increased pathogen systemic dissemination in the absence of complement activity. To test this, we measured *C. rodentium* WT and Δ*rfaL* burdens in the liver/spleen of B6 and C3^-/-^ juveniles at enteric infection peak (10 days-post-inoculation). In both mouse strains, the WT was able to translocate and colonize the liver and spleen (Fig. 2E,F). In contrast, Δ*rfaL* was unable to establish systemic infection in B6 mice, with CFU recovered from liver/spleen at or below the limit of detection. In contrast, Δ*rfaL* was able to colonize the liver/spleen at similar rates than the WT in C3^-/-^ mice (Fig. 2E,F), suggesting O-ag plays an essential role in natural, ID-associated bacteremia establishment by protecting *C. rodentium* from complement-mediated killing *in vivo*. Consistent with this, significantly higher C3b deposition was detected on Δ*rfaL* than WT recovered from liver and spleen of IP-injected B6 juveniles (Fig. S4).

**Figure 2.**
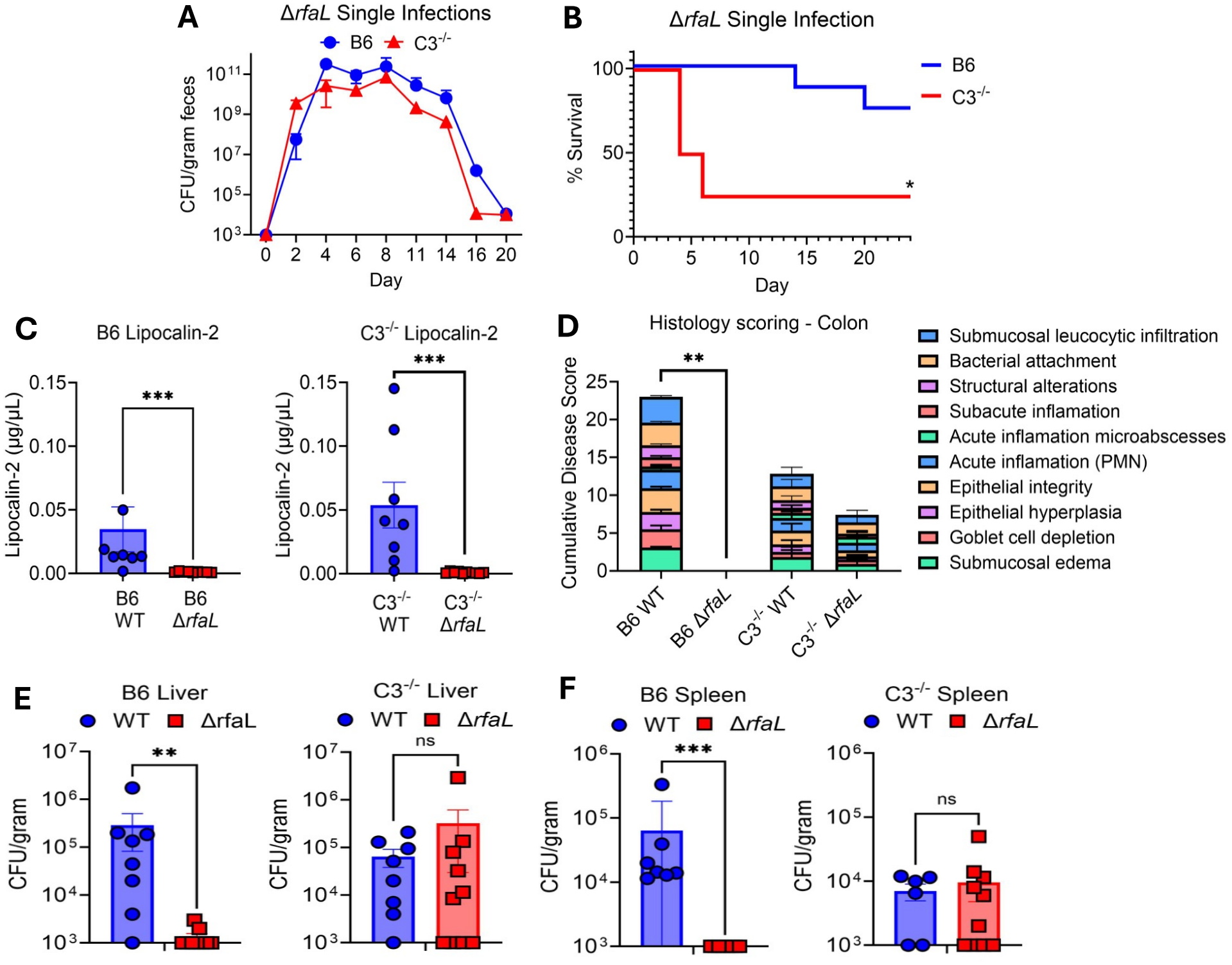
O-ag-deficient *C. rodentium* displays similar gut colonization, but differential mortality and systemic dissemination, in immunocompetent versus complement-deficient mice. **A –** Δ*rfaL* single infection in B6 (WT) and C3-KO (C3^-/-^) juveniles. CFU in feces during infection are shown. Δ*rfaL* displayed no significant differences in colonization between mouse strains. **B –** Survival percentages of B6 and C3-KO juveniles over the course of Δ*rfaL* single infection. C3-KO juveniles infected with Δ*rfaL* displayed marked mortality compared to B6 juveniles. **(C-F) –** B6 and C3-KO juvenile mice were infected with *C. rodentium* WT or Δ*rfaL*. 10 days post-infection, stool, colon tissue, liver, and spleen were collected. **C –** Fecal Lipocalin-2 levels of B6 and C3-KO infected juveniles. B6 and C3-KO juveniles infected with Δ*rfaL* exhibited reduced Lipocalin-2 levels than mice infected with the WT. **D –** Cumulative histological scores of colons from B6 and C3-KO infected juveniles. Each box indicates a specific parameter described on the right. B6 and C3-KO juveniles infected with Δ*rfaL* exhibited reduced intestinal pathology than mice infected with the WT. Statistical analysis: Mann-Whitney U test (WT vs Δ*rfaL* comparisons). **E –** WT and Δ*rfaL* CFU in livers of B6 and C3-KO infected juveniles. B6 juveniles infected with *C. rodentium* WT displayed significantly higher CFU compared to B6 counterparts infected with Δ*rfaL*. There were no significant differences in liver CFU between bacterial strains in C3-KO mice. **F –** WT and Δ*rfaL* CFU in spleen samples of B6 and C3-KO infected juveniles. B6 juveniles infected with *C. rodentium* WT displayed significantly higher CFU compared to B6 counterparts infected with Δ*rfaL*. There were no significant differences in spleen CFU between bacterial strains in C3-KO mice. Data are shown as mean ± SEM (A, C-F). Statistical analyses: two-tailed unpaired T-test (A); Log-rank Mantel-Cox test (B); Mann-Whitney U test (C-F). **\*** p≤0.05 **\*\***p≤0.01 ***p≤0.001

## DISCUSSION

Prior *in vivo* studies have shown O-ag is a major immunogenic component of *C. rodentium*, eliciting pathogen-specific antibody responses during infection (20,21). Furthermore, in a hypersusceptible adult model of infection (C3H/HeJ mice), another O-ag-deficient *C. rodentium* mutant (Δ*wfaP*) successfully colonized the intact gut, albeit with delayed kinetics and mortality rates than the WT strain (20). Together with our results in juvenile B6 mice, these findings indicate that O-ag-deficient mutants are still capable of colonizing the gut of hypersusceptible hosts under normal conditions, with lethality outcomes likely shaped by underlying host genetics or immune deficiencies (e.g., C3H/HeJ genetic background, complement deficiency).

The results presented here also support our prior inference that O-ag is required for pathogen competition with the mature gut microbiota (13). Moreover, we uncover previously unrecognized age- and microbiota-dependent requirements for O-ag in the host. Specifically, we found that juvenile SPF mice and gnotobiotic adult mice harboring juvenile microbiotas are permissive to colonization by O-ag-deficient *C. rodentium*. In contrast, the O-ag mutant exhibits impaired colonization in SPF and gnotobiotic mice with adult microbiotas. Further, we demonstrate this effect is mediated, at least in part, by microbiota-derived metabolites because adult, but not juvenile, cell-free cecal extracts inhibit mutant growth. Identifying commensal-derived compounds responsible for restricting colonization by O-ag-deficient enteric pathogens represents an open avenue for future investigations. These studies may help identify protective commensals and metabolites for development of new treatments against ID caused by virulent and attenuated bacteria in highly vulnerable populations.

Although it is well-known O-ag is required for pathogen resistance to serum inhibitory factors, including complement (15,17,22,23), the mechanistic basis for this requirement has not been studied in detail in models of natural systemic translocation across the gut. Using our model of ID-associated bacteremia in juvenile mice, we show that O-ag-deficient *C. rodentium* can translocate across the gut and establish robust systemic infection in complement-deficient, but not complement-sufficient, hosts. This is clinically relevant as newborns have reduced levels of complement components (24,25) and O-ag-deficient pathogens have been identified among clinical isolates (12). In closing, our results demonstrate that O-ag-deficient enteric bacterial pathogens can retain the capacity to outcompete the early-life microbiota during intestinal colonization while remaining capable of systemic dissemination in hosts with impaired complement function. These findings may inform ongoing efforts to develop effective vaccines against EHEC and other bacterial pathogens (21,26), particularly for children and immunocompromised individuals.

## DATA AVAILABILITY

16S rRNA gene sequencing data are available at the NCBI Sequence Read Archive under BioProject No. PRJNA1559746. This paper does not report original code.

## ACKNOWLEDGMENTS

This research was supported by NIH grant R00AI159620 and UW-Madison Start-up funds to G.C.-F. The authors thank Annelise Resende and the University of Wisconsin-Gnotobiotic Shared Resource for technical support with experiments in GF/Gnotobiotic mice. We thank the Gumperz and Sperandio Labs in the Department of Medical Microbiology & Immunology at UW–Madison for sharing equipment for the execution of specific experiments in this study. Authors also thank Patricia Tran for advice on bioinformatic analysis.

## AUTHOR CONTRIBUTIONS

Aidan Schmidt conceived and designed the project, performed most experiments, prepared final figures, and helped write the manuscript; Erick D. Nilsen performed experiments, prepared figure panels, and helped write the manuscript; Marienela Y. Heredia, performed flow cytometry assays; Victoria G. Zalutskiy performed bioinformatic analyses; Andres Mejia, performed histological analyses; Gustavo Caballero-Flores conceived, designed, and supervised the project – reviewed, edited, and finalized the manuscript with contributions from all authors.

